# Perimenstrual Exacerbation of Symptoms in Borderline Personality Disorder: Evidence from Multilevel Models and the Carolina Premenstrual Assessment Scoring System

**DOI:** 10.1101/192658

**Authors:** Tory A. Eisenlohr-Moul, Katja M. Schmalenberger, Sarah A. Owens, Jessica R. Peters, Danyelle N. Dawson, Susan S. Girdler

## Abstract

**Background:** Individuals with borderline personality disorder (BPD) suffer from a constellation of rapidly shifting emotional, interpersonal, and behavioral symptoms. The menstrual cycle may contribute to symptom instability among females with this disorder.

**Methods:** Fifteen healthy, unmedicated females with BPD and without dysmenorrhea reported daily symptoms across 35 days. Urine luteinizing hormone (LH) and salivary progesterone (P4) were used to confirm ovulation and cycle phase. Cyclical worsening of symptoms was evaluated using (1) phase contrasts in multilevel models and (2) the Carolina Premenstrual Assessment Scoring System (C-PASS; Eisenlohr-Moul et al., 2017b), a protocol for evaluating clinically significant cycle effects on symptoms.

**Results:** Most symptoms demonstrated midluteal worsening, a perimenstrual peak, and resolution of symptoms in the follicular or ovulatory phase. Post-hoc correlations with person-centered progesterone revealed negative correlations with most symptoms. Depressive symptoms showed an unexpected delayed pattern in which baseline levels of symptoms were observed in the ovulatory and midluteal phases, and exacerbations were observed during both the perimenstrual and follicular phases. The majority of participants met C-PASS criteria for clinically significant (>=30%) symptom exacerbation. All participants met the emotional instability criterion of BPD, and no participant met DSM-5 criteria for premenstrual dysphoric disorder (PMDD).

**Conclusions:** Females with BPD may be at elevated risk for perimenstrual worsening of emotional symptoms. Longitudinal studies with fine-grained hormonal measurement as well as hormonal experiments are needed to determine the pathophysiology of perimenstrual exacerbation in BPD.

---

Individuals with borderline personality disorder (BPD) suffer from a chronic and disabling combination of emotional, cognitive, and behavioral psychiatric symptoms. Diagnostic criteria for BPD (five of which must be met for diagnosis) highlight instability and flux in various domains, including unstable moods, reactive and uncontrolled anger (exacerbated by a tendency to ruminate on angry experiences (Baer & Sauer, 2011; Peters et al, 2017), changing identities, frequent feelings of emptiness, turbulence in interpersonal relationships, fear of abandonment, a tendency to become paranoid or dissociate during times of stress, and damaging, impulsive behaviors. Recurrent, self-injurious thoughts and behaviors are also a symptom, and roughly 10% of people with BPD die from suicide (Paris & Zweig-Frank 2001). Individuals with BPD are heavy utilizers of psychiatric treatment including therapy, hospitalizations, and pharmacotherapy (Bender et al, 2001). Treatment for BPD can be beneficial, however, it is often extremely challenging (Aviram et al, 2006), and quality of life following treatment often remains low despite improved functioning (Linehan *et al*. 2006).

Research on the biology of BPD has generally focused on stable risk factors, whereas relatively little research examines the proximal causes of *instability* in BPD symptoms. Given that the ovarian steroid hormones estradiol (E2) and progesterone (P4) regulate nearly every brain system implicated in emotional disorders (Schiller *et al*. 2016), the rapid changes in ovarian steroid hormones that occur across the monthly female menstrual cycle could represent one source of fluctuating biological vulnerability to BPD symptoms. The present study uses a longitudinal design and standardized diagnostic system to describe the impact of menstrual cycle phase on symptoms in unmedicated females with BPD.

The prototypical menstrual cycle is 28 days and has two parts: the *follicular phase* from menses to ovulation (low P4, increasing E2 that peaks prior to ovulation), and the *luteal phase* from ovulation to the next menses (high P4, secondary slower and generally lower peak in E2). E2 and P4 fall precipitously prior to and during the first few days of menses (described as perimenstrual steroid withdrawal), and the cycle begins again. Among patients with the primary cyclical mood disorder of DSM-5 premenstrual dysphoric disorder (PMDD), symptoms peak in the perimenstrual window (Epperson *et al*. 2012, Hartlage *et al*. 2002), during perimenstrual steroid withdrawal. This dovetails with preclinical findings that ovarian steroid withdrawal precipitates anxious and depressive phenotypes (Stoffel & Craft 2004; Smith *et al*. 2006).

Although most females do not show clinically significant menstrual cycle effects on emotional (Schwartz *et al*. 2012; Ben Dor *et al*. 2013; Hengartner *et al*. 2017) or cognitive (Schmidt *et al*. 2013; Leeners *et al*. 2017) symptoms, experiments demonstrate that a subset of females suffer from *abnormal sensitivity to normal ovarian steroid changes*. This ***hormone sensitivity*** manifests as emotional, cognitive, and behavioral changes appearing only or primarily in the context of normative ovarian steroid changes, such as the midluteal and perimenstrual cycle phases, as in PMDD (Schmidt *et al*. 1998) or perimenstrual exacerbation (PME) of symptoms related to a chronic underlying disorder; during pregnancy and postpartum (Bloch *et al*. 2000); and during the menopause transition (Schmidt *et al*. 2015; Gordon *et al*. 2016). The content of emotional symptoms most commonly observed in these *reproductive mood disorders* overlaps to a greater degree with BPD than with major depressive disorder: Reproductive mood disorders are often characterized by *high arousal negative affect* and *interpersonal* symptoms like anger, irritability, and interpersonal sensitivity, whereas depressive symptoms are less common (Freeman *et al*. 2011; Eisenlohr-Moul *et al*. 2017a; Putnam *et al*. 2017). While being female with a mood or anxiety disorder may not in itself equate to a high risk of PME (Stein *et al*. 1989; Hartlage *et al*. 2004), PME may be more prevalent in a disorder such as BPD already characterized by instability of these symptoms.

Little prospective evidence is available to evaluate the hypothesis that females with BPD are at high risk of PME. Only one study has examined the impact of the menstrual cycle in patients with BPD (Ziv, 1995; N=14). This study reported no evidence of reliable symptom changes from the premenstrual to postmenstrual phases of the cycle. However, participants took psychotropic medications, and ovulation was not confirmed; the latter point is critical, since only ovulatory cycles are symptomatic in PMDD or PME (Hammarbäck *et al*. 1991). Another series of studies (De Soto *et al*. 2003) found that undergraduate females with high BPD features at baseline showed greater increases in symptoms after beginning oral contraceptives (OCs; N=46), and during cyclical steroid changes (N=57), both suggesting risk of hormone sensitivity in BPD. In a final study of undergraduate females blinded to the purpose of the study (*N*=40), individuals with elevated trait BPD features showed their highest BPD symptoms when P4 was higher-than-average and E2 was lower-than-average, a hormonal profile found primarily in the midluteal and perimenstrual phases (Eisenlohr-Moul *et al*. 2015). These findings converge to suggest the possibility of heightened risk for psychiatric hormone sensitivity and related PME in BPD.

The present study examines the prospective impact of cycle phase (in ovulatory, normal menstrual cycles) on daily self-reported symptoms in community-recruited, unmedicated females with BPD (*N* = 15) *not* recruited for perceived perimenstrual symptoms. Although participants collected saliva eight times to verify cycle phase, the present study was not powered to examine hormonal effects, instead focusing on cycle phase. In addition to examining the significance and size phase differences, we utilize a standardized diagnostic system, the Carolina Premenstrual Assessment Scoring System (C-PASS; Eisenlohr-Moul *et al*. 2017b), to describe, in absolute terms, the clinical significance of PME in each participant.

## Hypotheses

We tested the following hypotheses:

1. (1)**<u>Perimenstrual Exacerbation (Relative to all other Phases)</u>**: We predicted that participants with BPD would also show their greatest emotional symptom exacerbation during the <u>perimenstrual</u> phase relative to each other phase comparable to the symptom emergence pattern in PMDD.
2. (2)**<u>Midluteal Exacerbation (Relative to Follicular and Ovulatory Phases)</u>**: Given that women with primary cyclical mood disorder (i.e., PMDD) generally *begin* to have emotional symptoms during the midluteal phase (when P4 rises and begins to fluctuate), we also predicted that participants with BPD would show symptom exacerbation in the <u>midluteal</u> phase relative to follicular and ovulatory phases, but that midluteal symptoms would still be lower than the perimenstrual phase.
3. (3)**<u>>50% Sample Diagnosis with C-PASS Premenstrual Symptom Elevation</u>**: Given overlapping risk factors and similar core symptom content in individuals with primary cyclical mood disorder (i.e., PMDD) and individuals with BPD, we predicted that a majority (>50%) of this BPD sample would meet the C-PASS diagnostic criterion of *premenstrual elevation* (worsening of at least 30% premenstrually vs. postmenstrually) on at least one emotional symptom. However, given that BPD is a chronic emotional disorder that does not remit fully for weeks at a time (as is required in PMDD), we predicted that no participant would meet DSM-5 criteria for PMDD in the examined cycle.

## Methods

### Procedure

We recruited participants using listservs, flyers, and social media advertisements seeking “women with emotional and interpersonal problems that interfere with life”. These materials *did not refer to the menstrual cycle* to avoid self-selection. Interested individuals were directed to an online screening, including the *Personality Diagnostic Questionnaire—Borderline Personality Disorder Subscale* (PDQ; Hyler *et al*. 1992) and eligibility questions. Exclusionary criteria were: dysmenorrhea in the past 3 months, current pregnancy or breastfeeding, psychotropic medication or OC use, illicit drug use, and chronic medical conditions, and schizophrenia or manic episodes. Menstrual cycles between 25-35 days were required.

We contacted Individuals for phone screening if they met inclusion criteria and had a PDQ score of 5 out of 9 BPD criteria. The phone screen included an interview version of the PDQ. We invited those providing symptom examples indicating significant “pathology, persistence, and pervasiveness” (First *et al*. 1997) on at least 5 of 9 BPD symptoms for an enrollment visit. This visit included a SCID-PD (BPD only) interview with a clinical psychologist. If they met BPD criteria, they completed questionnaires and oriented to study procedures (daily survey completion, urine ovulation testing, and salivary collection). Participants completed daily surveys for 35 days starting on their next menstrual period start day, daily urine ovulation testing based on cycle length, and passive drool saliva samples on 8 days across the cycle. Prospective charting of daily symptoms was critical due to the well-documented lack of correspondence between cross-sectional retrospective reports of premenstrual symptoms and prospective patterns of symptom change (e.g., Roy-Byrne et al., 1986). Participants received $15 for the enrollment visit, and $100 for the larger study. All procedures were approved by the university institutional review board. Data were collected 2015-2016.

### Participants

310 individuals completed online screening; 112 were eligible for phone screening (exclusion due to insufficient PDQ severity (*n*=143), hormone use (*n*=42), irregular cycles (*n*=6), or medical conditions (*n*=7)). Of those, 43 completed the phone screen; we invited 31 of those for an enrollment visit (exclusion due to subthreshold PDQ examples (*n*=8), hormone use (*n*=2), pregnancy/nursing (*n*=2)). Of those, 18 completed the initial visit. Seventeen met SCID-BPD criteria and enrolled. One participant was noncompliant and one did not ovulate; the final sample included 15 medically healthy, unmedicated females with BPD. Demographics and clinical characteristics are provided in Table 1. All participants reported a lifetime history of psychotherapy; 1 was currently in psychotherapy.

**Table 1.** Sample Descriptive Information (*N* = 15)

| Variable | Mean (SD); n (%) | Range |
| --- | --- | --- |
| **General Characteristics** |  |  |
| Age | 32.16 (8.64) | 19-44 |
| African American Race | 5 (33%) | - |
| Hispanic Ethnicity | 1 (6%) | - |
| **Education Level** |  |  |
| Trade School | 1 (6%) | - |
| Some College | 7 (47%) | - |
| College Grad | 7 (47%) | - |
| **Marital Status** |  |  |
| Married | 2 (15%) | - |
| Living with Partner | 5 (33%) | - |
| Divorced | 3 (20%) | - |
| Never Married | 5 (33%) | - |
| Average Cycle Length in Days | 28.97 (1.32) | 26-35 |
| Age at Menarche in Years | 11.24 (1.08) | 10-14 |
| % with Biological Children | 4 (26%) | - |
| **Clinical Characteristics** |  |  |
| **Trauma Exposure** |  |  |
| Childhood Trauma Questionnaire (CTQ) | 43.00 (12.08) | 28-64 |
| % Significant Childhood Physical Abuse ($\geq 8$) | 7 (47%) | - |
| % Significant Childhood Sexual Abuse ($\geq 8$) | 4 (27%) | - |
| % Significant Childhood Physical Neglect ($\geq 8$) | 7 (47%) | - |
| % Significant Childhood Emotional Abuse ($\geq 10$) | 7 (47%) | - |
| % Significant Childhood Emotional Neglect ($\geq 15$) | 5 (33%) | - |
| **Borderline Symptom Assessments** |  |  |
| PAI-BOR Score | 44.23 (9.84) | 29-53 |
| Number of SCID-BPD Criteria Met | 6.89 (1.26) | 5-8 |
| % Meeting Fear of Abandonment Criterion | 10 (67%) | - |
| % Meeting Affective Lability Criterion | 15 (100%) | - |
| % Meeting Identity Disturbance Criterion | 8 (53%) | - |
| % Meeting Interpersonal Disturbances Criterion | 14 (93%) | - |
| % Meeting Emptiness Criterion | 11 (73%) | - |
| % Meeting Anger Criterion | 10 (67%) | - |
| % Meeting Paranoia/Dissociation Criterion | 8 (53%) | - |
| % Meeting Self-Harm Criterion | 6 (40%) | - |
| % Meeting Impulsivity Criterion | 11 (73%) | - |
| **Comorbidity Assessments** |  |  |
| Depression (BDI) | 13.66 (4.37) | 2-29 |
| % Minimal (0-13) | 5 (33%) | - |
| % Mild (14-19) | 5 (33%) | - |
| % Moderate (20-28) | 3 (30%) | - |
| % Severe (29-63) | 2 (13%) | - |
| Anxiety (STAI-T) | 45.36 (8.86) | 29-56 |

Note. CTQ = Childhood Trauma Questionnaire. STAI=State-Trait Anxiety Inventory (Trait Form). BDI = Beck Depression Inventory. PAI-BOR = Personality Assessment Inventory – Borderline Subscale. SCID-BPD = Structured Clinical Interview for Personality Disorders – Borderline Personality Disorder Module.

### Baseline Measures

In addition to the SCID-II for BPD, participants completed several questionnaires during their baseline visit: a) demographics, b) the borderline subscale of the *Personality Assessment Inventory* (PAI-BOR; Morey 2014), c) the *Childhood Trauma Questionnaire* (CTQ; Bernstein & Fink 1998), d) the *Beck Depression Inventory* (BDI; Beck & Steer 1987) and the trait form of the *State-Trait Anxiety Inventory* (STAI; Spielberger 1983), and e) items from the *Premenstrual Assessment Form* (Halbreich *et al*. 1982) to assess perceptions of menstrually-related symptom change. Participants were asked to rate their typical degree of premenstrual symptom exacerbation for depressed mood, hopelessness, worthlessness or guilt, anxiety, anger, irritability, and interpersonal conflicts on a scale where 1=no change, 2=mild change, 3=moderate change, and 4=severe change.

### Daily Measures

Participants completed daily questionnaires via online survey; links were sent daily via text message and email.

#### Daily Record of Severity of Problems (DRSP)

The DRSP (Endicott *et al*. 2006) is a daily measure developed to reliably detect menstrually-related changes (e.g. PMDD, PME) in psychiatric and physical symptoms (<u>depression, anxiety, anhedonia, hopelessness, emotional overwhelm, rejection sensitivity, anger/irritability, interpersonal conflict</u>, and <u>physical symptoms</u> (average of breast tenderness, breast swelling, bloating, joint or muscle pain)). Items are rated from 1=Absent to 5=Extreme.

#### PME Diagnosis Using the Carolina Premenstrual Assessment Scoring System (C-PASS) for the DRSP

The degree of premenstrual symptom elevation (and associated clinical significance cutoffs) can be determined with the C-PASS, a standardized scoring system <u>used in conjunction with daily DRSP data</u> to make the PMDD or PME diagnosis. The C-PASS operationalizes the DSM-5 PMDD diagnosis into diagnostic dimensions and numerical diagnostic thresholds validated against expert diagnosis based on daily ratings (Eisenlohr-Moul *et al*. 2017b). Here, the C-PASS provides standardized criteria for identifying whether or not a participant’s daily ratings demonstrate clinically significant PME (>= 30% increase premenstrually vs. postmenstrually) of any emotional symptom.

#### Additional BPD Symptoms

Three additional BPD-relevant symptoms were measured daily: <u>shame</u> was measured using two items from the shame subscale of the State Shame and Guilt Scale (Marschall et al. 1994); “I wanted to sink into the floor and disappear”, and “I felt humiliated, disgraced”). Daily <u>felt invalidation</u> was measured with two items created for the purposes of this study (“I felt that someone wanted to me to stop feeling the way I was feeling”, “I felt that someone didn’t take my feelings seriously”). Daily <u>anger rumination</u> was measured with two items from the Anger Rumination Scale (Sukhodolsky et al. 2001); “I kept thinking about events that have angered me”, and “I turned anger-provoking situations over and over again in my mind”). Each of these items were rated from 1 (Not at All) to 5 (Extremely).

### Ovulation Testing and Salivary Ovarian Steroids

Urine luteinizing hormone tests (LH; Clearblue ovulation testing kit) confirmed ovulation. This test has been validated against ultrasound and has a 40 pg/mL LH threshold, reducing false positives. Ovulation can occur 12-36 hours after the LH surge; here, the day after the positive test was considered ovulation. Testing started 7 days after menses onset until a positive result. Participants provided saliva, assayed for E2 and P4, on 8 days: days 2, 3, 8, and 9 following menstrual onset and days 2, 3, 8, and 9 following positive LH surge test. This method was used to increase sample reliability for cycle phase validation. Participants collected 5 mL of saliva in polypropylene vials (passive drool) each afternoon 4-6pm, froze samples in their home freezer in a temperature stability pouch, and returned them to the lab after study completion. Afternoon sampling was chosen to increase participant compliance based on poor morning compliance in our pilot data. Hormones were assayed using RIA kits purchased from Salimetrics (Carlsbad, CA) at the UNC Chapel Hill School of Medicine Core Laboratory.

#### Reliability and Validity of Salivary E2 and P4

Intra-assay CVs were 6.3% and 4.0% for E2 and P4, respectively. Interassay CVs were 8.9% and 5.5% for E2 and P4, respectively. Per Salimetrics validation reports, the sensitivity of the Salimetrics RIA method is .1pg/ml for E2, and 5pg/ml for P4, and salivary E2 and P4 should show similar cyclical changes to serum E2 and P4 across the cycle. In our sample, P4 demonstrated the expected post-ovulatory rise in each participant. However, E2 demonstrated virtually no cyclical change for any participant. Given that ovulation was confirmed and P4 showed expected patterns, this lack of E2 clearly indicated sampling, storage, or assay issues. We suspect that at-home storage for one month, combined with the diurnal pattern reducing afternoon E2, may have caused this issue. Therefore, we proceeded to use the P4 values, and did not use the E2 values.

### Menstrual Cycle Phase Coding

Determination of cycle phases—*ovulatory, midluteal*, *perimenstrual*, and *follicular*—was based the following: (1) forward and backward count (Edler *et al*. 2007), (2) day of LH surge, and (3) exponential P4 rise following ovulation. First, the ovulatory phase was determined using backward count (days -15 to -12), and was validated using the LH surge ovulation test and a substantial postovulatory rise in P4. Next, the perimenstrual phase was determined using backward and forward count (days -4 to +3, where day +1 is menses onset and there is no day 0), and was validated by time since positive LH surge (14+/-2 days). The midluteal phase was defined as the days falling between the ovulatory and perimenstrual phases, and was validated by elevated P4. The follicular phase was defined as days falling between the perimenstrual and ovulatory phases, and was validated by low P4.

### Data Analysis and Power

To improve interpretability and isolate the within-person component of the outcome, each outcome was person-standardized (as in Edler *et al*. 2007; Klump *et al*. 2013). To test the first three predictions, multilevel models evaluated the impact of cycle phase (coded as a categorical variable, with alternating reference phases). SAS PROC MIXED accounted for the nested structure of observations (i.e., random intercept) as well as the autocorrelation of today’s rating with yesterday’s rating. Random effects of cycle phase contrasts were not included as they did not improve model fit; this indicates that the impact of the menstrual cycle on symptoms were roughly uniform in this sample. Five-day rolling averages of each person-standardized outcome were utilized for graphical depictions.

To determine power for phase contrasts, we calculated the design effect to determine the smallest detectible effect size (Snijders 2005). Each participant provided up to 35 daily surveys (lower-level *N* was 490), and symptoms generally showed low intraclass correlations (i.e., a high degree of flux within a person, average ICC for DRSP items = .18). Power for these models was adequate to detect conventionally small-to-medium sized contrasts between phases (average smallest detectible effect size was an *f^2^* of .10). That is, the study was powered to detect significant phase contrasts accounting for at least 10% of the variance in the daily outcome. Age was considered as a covariate but ultimately omitted as it was not associated with outcomes and did not alter our pattern of findings.

To test hypothesis 4, we evaluated whether each participant met the C-PASS diagnostic criterion of *premenstrual elevation* (over the postmenstrual baseline) on at least one emotional symptom (Eisenlohr-Moul *et al*. 2017b). In contrast to PMDD, which requires luteal phase elevation *and* full remission of symptoms in the postmenstrual week, clinically significant PME can be diagnosed when at least one emotional symptom demonstrates a >=30% *premenstrual* week elevation of symptoms relative to the postmenstrual week (with no requirement of a remission week as in PMDD; see Eisenlohr-Moul et al., 2017b for calculation details). This 30% change threshold for clinically significance was delineated by the NIMH work group for PMDD.

## Results

### Descriptives

Table 1 provides demographic and clinical characteristics. The sample was diverse with regard to age and race. Participants reported high BPD symptoms on the PAI-BOR, moderate depression on the BDI, moderate anxiety on the STAI, and significant experiences of childhood trauma; these characteristics are consistent with typical comorbidities and childhood experiences in BPD. Participants cross-sectionally reported an average perception of *mild* PME of emotional symptoms. 93% of surveys were completed, and 90% of salivary samples were completed.

### Contrasting BPD Symptom Severity Across Menstrual Cycle Phases

Results of multilevel models testing phase effects on symptoms are presented in Table 2. All symptoms demonstrated strong cyclical changes; Figures 1-3 depict patterns of symptom change across cycle day and cycle phase for several exemplar symptoms. For each model reported below, additional robustness-check models controlled for person-standardized physical symptoms; we observed no substantive model changes, suggesting PME is not an epiphenomenon of perimenstrual physical pain (e.g., dysmenorrhea). Graphs by day and cycle phase are provided for each outcome in Supplement 1. Graphs include smoothed, person-standardized P4 values.

**Table 2.** *Multilevel Model Results: Within-Person Menstrual Cycle Phase Contrasts for Intrapersonal and Interpersonal Symptoms*

|  | **Within-Person Phase Contrasts (Standardized Fixed Effects)** |  |  |  |  |  |
| --- | --- | --- | --- | --- | --- | --- |
|  | Perimenstrual Phase<br>(Falling E/P) |  |  | Midluteal Phase<br>(High Stable E/P) |  | Follicular Phase<br>(Low Stable E/P) |
|  | v. Follicular<br>(Low Stable E/P) | v. Midluteal<br>(High Stable E/P) | v. Ovulatory<br>(Rising E/Low P) | v. Ovulatory<br>(Rising E/Low P) | v. Follicular<br>(Low E/P) | v. Ovulatory<br>(Rising E/Low P) |
| **Core Cyclical Emotional Symptoms (DRSP)** |  |  |  |  |  |  |
| Depression | -.12 (.35) | -.51*** (.14) | -.51** (.17) | -.01 (.17) | .39** (.14) | -.39** (.15) |
| Hopelessness | -.12 (.12) | -.44** (.14) | -.71*** (.16) | -.26 (.17) | .32* (.13) | -.59*** (.16) |
| Anxiety | -.61*** (.13) | -.34* (.15) | -.70*** (.17) | -.37* (.18) | -.28 (.14) | -.09 (.17) |
| Rejection Sens | -.26* (.12) | -.33* (.14) | -.40* (.16) | -.07 (.17) | .07 (.13) | -.13 (.15) |
| Anger/Irritability | -.51*** (.13) | -.24* (.11) | -.76*** (.17) | -.52** (.18) | -.27* (.13) | -.24 (.17) |
| Interpersonal Conflict | -.71*** (.13) | -.33* (.14) | -.77*** (.17) | -.44* (.18) | -.37* (.14) | -.07 (.16) |
| **Additional Symptoms** |  |  |  |  |  |  |
| Anhedonia | -.12 (.13) | -.35* (.15) | -.34* (.17) | .004 (.19) | .22 (.14) | -.22 (.17) |
| Overwhelmed, Can't Cope | -.50*** (.12) | -.36* (.14) | -.70*** (.17) | -.35 (.16) | .14 (.13) | .21 (.16) |
| Shame | -.22 (.12) | -.28* (.12) | -.54** (.16) | -.26 (.17) | .49*** (.13) | -.76*** (.16) |
| Felt Invalidation | -.60*** (.13) | -.45** (.15) | -.53** (.17) | -.07 (.18) | -.15 (.14) | -.08 (.18) |
| Anger Rumination | -.54*** (.13) | -.42** (.15) | -.50** (.17) | -.07 (.18) | -.12 (.14) | .04 (.17) |
| Physical Symptoms | -.36** (.13) | -.39* (.15) | -.79*** (.17) | -.41* (.18) | -.01 (.14) | -.43* (.17) |

*Note.* \* $p < .05$ ; \*\* $p < .01$ \*\*\* $p < .001$ . Hormone profile labels for the cycle phases are descriptive of typical cycles, since E was not assessed and P was assessed in only some of the cycle phases. For presentation in this table only, outcomes have been person-standardized so that estimates represent the number of standard deviations from one's mean (0); therefore, estimates reflect the average number of personal standard deviations by which the two phases differ. DRSP=Daily Record of Severity of Problems.

**Figure 1.**
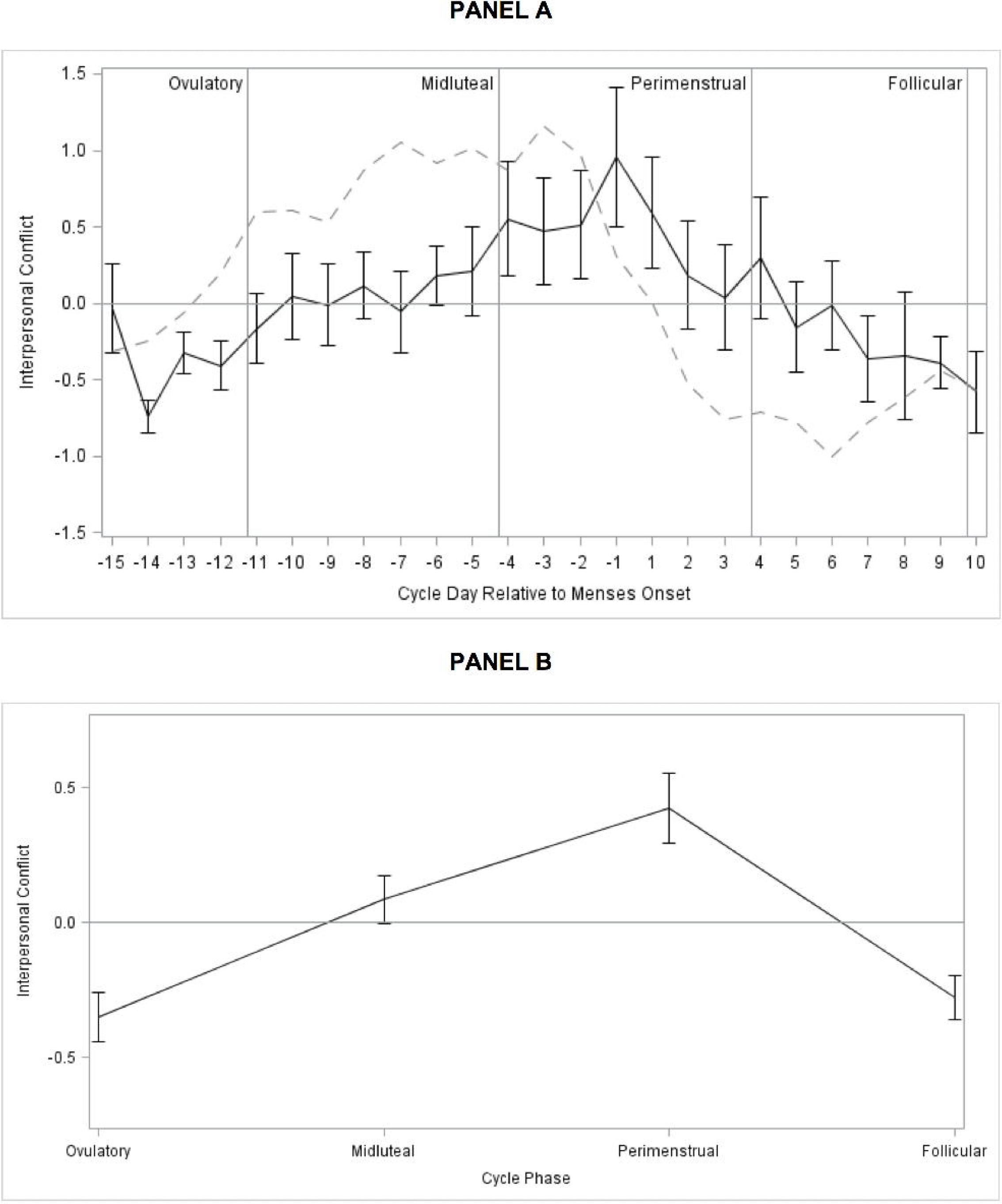
Person-Standardized Progesterone (dashed line) and Interpersonal Conflict Across Menstrual Cycle Day (Panel A) and Person-Standardized Interpersonal Conflict Across Cycle Phase (Panel B) in 15 People with BPD.

**Figure 2.**
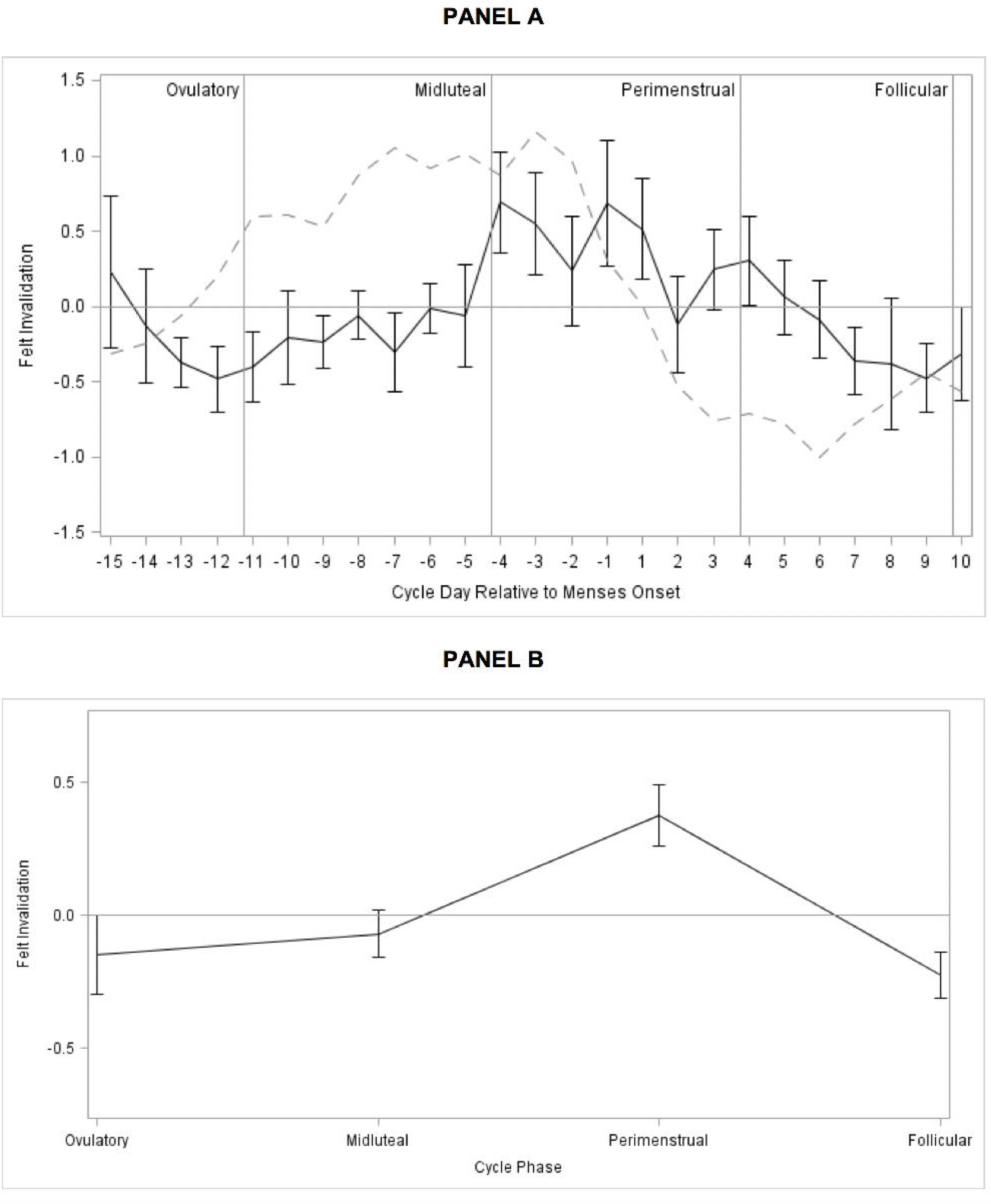
Person-Standardized Progesterone (dashed line) and Felt Invalidation Across Menstrual Cycle Day (Panel A) and Person-Standardized Felt Invalidation Across Cycle Phase (Panel B) in 15 People with BPD.

**Figure 3.**
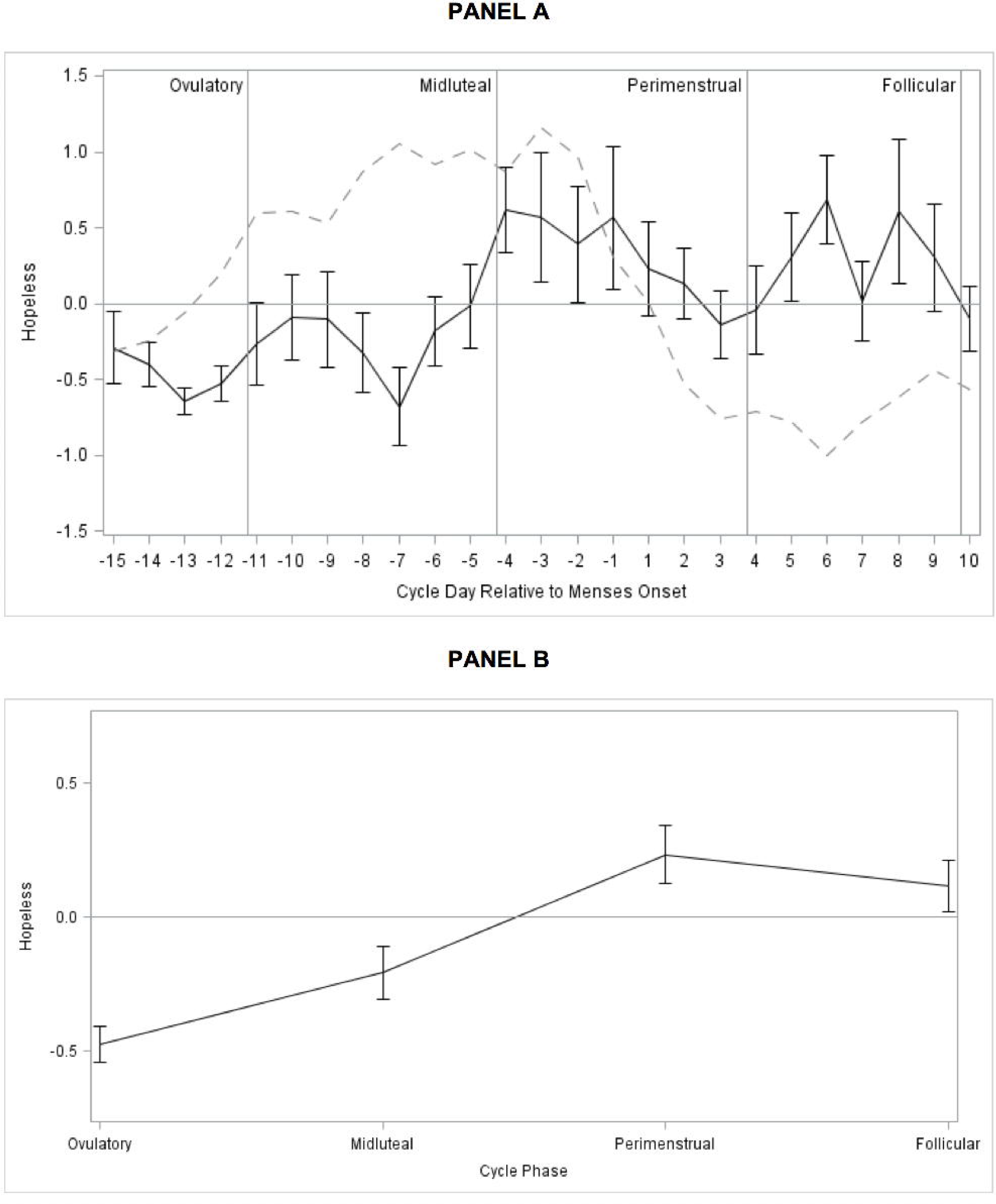
Person-Standardized Progesterone (dashed line) and Hopelessness Across Menstrual Cycle Day (Panel A) and Person-Standardized Hopelessness Across Cycle Phase (Panel B) in 15 People with BPD.

### Testing Hypothesis 1: Perimenstrual Exacerbation (vs. all other Phases)

We predicted that the perimenstrual phase would be associated with greater symptoms relative to each other cycle phase. Symptoms were greater in the perimenstrual phase than in the **midluteal** and **ovulatory** phases for all symptoms. Symptoms were also greater in the perimenstrual phase than in the **follicular** phase for the majority of symptoms, although no differences were observed between the perimenstrual and follicular phases for depression, hopelessness, anhedonia, or shame, indicating that the exacerbation of those four symptoms, which began perimenstrually, *continued on into the* follicular phase (see section for “Testing Hypothesis 3” below).

### Testing Hypothesis 2: Midluteal Exacerbation (vs. Follicular and Ovulatory Phases)

We predicted that participants would begin to show greater emotional symptoms in the midluteal phase relative to the follicular and ovulatory phases. Relative to the **ovulatory** phase, the <u>midluteal</u> phase was associated with worsening of four symptoms: anxiety, anger/irritability, interpersonal conflict, and physical symptoms. Relative to the **follicular** phase, the midluteal phase showed a mixed pattern of contrasts, with anger/interpersonal conflict symptoms showing expected midluteal worsening relative to the follicular phase (anger/irritability, interpersonal conflict), and several depressive symptoms showing an unexpected worsening in the follicular phase relative to the midluteal phase (depression, hopelessness, and shame).

### Exploratory Contrasts Between the Follicular and Ovulatory Phases

In general, the ovulatory phase demonstrated the lowest symptoms. Exploratory contrasts indicated that, relative to the follicular phase, the ovulatory phase showed lower levels of depression, hopelessness, shame, and physical symptoms, indicating that the perimenstrualto-follicular exacerbation of depressive symptoms observed above did return to the baseline level of symptoms in the ovulatory phase.

### Testing Hypothesis 4: >50% Sample Diagnosis with C-PASS Premenstrual Elevation

Given overlapping risk factors and core symptom content in PMDD and BPD, we predicted a majority (>50%) of the sample would meet the C-PASS diagnostic criterion of *premenstrual elevation* (>= 30% higher premenstrually vs. postmenstrually) on at least one emotional symptom. Results of individual C-PASS scoring of daily DRSP ratings revealed 11 of 15 participants (73%) showed clinically significant PME (>= 30% premenstrual symptom elevation relative to the postmenstrual week) of at least one DRSP emotional symptom.

Exclusion of the first three days of menses may have omitted the most symptomatic days (Pearlstein *et al*. 2007); therefore, we also conducted analyses replacing the pre- and post-menstrual means with perimenstrual and follicular means (as defined in analyses for Hypotheses 1-2)), respectively; this did not alter the 73% outcome. Given the observed follicular exacerbation of depressive symptoms, we also conducted analyses replacing the pre- and post-menstrual means with those from the perimenstrual and *ovulatory* phases, respectively. Fourteen out of fifteen participants (93%) showed clinically significant change of at least one emotional symptom from the ovulatory to the perimenstrual phase.

As predicted, no participant in the present study met C-PASS PMDD criteria for any symptom in the examined cycle. Since all participants continued to have moderate-or-greater symptoms in the postmenstrual week (consistent with BPD), they failed to meet the PMDD diagnostic criterion of *absolute clearance* (Eisenlohr-Moul et al., 2017b).

### Post-Hoc Associations of P4 with Daily Symptoms

Despite low statistical power, we examined preliminary within-person correlations between P4 and symptoms. Consistent with preclinical evidence that steroid withdrawal elicits relevant phenotypes (Stoffel and Craft, 2004; Smith et al., 2006), we observed negative correlations of person-centered P4 with a variety of symptoms: depression (within-person correlation of P4 with depression on the same day, γ=-.0021, *p*=.042), hopelessness (γ=-.0015, *p*=.039), anxiety (γ=-.0056, *p*=.012), and rejection sensitivity (γ=-.0037, *p*=.027), anger/irritability (γ=-.0021, *p*=.042), anhedonia (γ=-.0031, *p*=.044), overwhelm (γ=-.0045, *p*=.011), felt invalidation (γ=-.005, *p*=.022), and physical symptoms (γ=-.0033, *p*=.018). Upon inspection of individual symptom plots, these correlations appeared to be driven by heightened symptoms during perimenstrual and follicular withdrawal from P4.

## Discussion

We sought to describe menstrual cycle patterns of symptom exacerbation in unmedicated females with BPD. Despite reporting only mild perceived PME in the baseline survey, daily ratings unequivocally demonstrated a high risk of perimenstrual symptom exacerbation in all symptoms. The observed symptom patterns can be described as follows. First, all symptoms generally show lowest levels in the ovulatory phase. Second, in the midluteal phase, high-arousal symptoms worsen, but are not yet accompanied by exacerbation of other symptoms. Third, in the perimenstrual phase, all symptoms become exacerbated, with further exacerbation of high-arousal symptoms, but also increased experience of physical discomfort and feelings of invalidation, shame, rejection sensitivity, anger rumination, emotional overwhelm, depression, anhedonia, and hopelessness. Finally, although high-arousal symptoms returning to baseline in the follicular phase, the lower-arousal depressive symptoms continued to be exacerbated in the follicular phase before returning to baseline at ovulation and the midluteal phase. Critically, physical symptoms did not account for PME of emotional symptoms. C-PASS diagnostic procedures indicated clinically significant PME in at least 11 of the 15 participants.

Although it is premature to comment on biological mechanisms of PME in BPD, we can compare and contrast observed patterns with those documented in PMDD, a disorder for which pathophysiology is better understood. Given their identical timing, PME of high-arousal symptoms in BPD may have a shared pathophysiology with PMDD—that is, a dysphoric response to luteal fluctuations in the GABAergic neurosteroid metabolites of P4 (Martinez *et al*. 2013; Schiller *et al*. 2014). However, regarding PME of depressive symptoms, the continued exacerbation in the follicular phase is not consistent with the follicular clearance observed in PMDD, suggesting a divergent pathophysiology. Females with BPD may have *additional* sensitivity to follicular withdrawal from—or low levels of—ovarian steroids. This withdrawal hypothesis receives preliminary support from our post-hoc finding of a negative within-person correlation between P4 and symptoms; howevere, these correlations should be interpreted with extreme caution due to low power. Finally, some have also postulated that females with BPD may also have greater sensitivity of serotonergic systems to E2 fluctuations (De Soto, 2007).

Several lines of research indicate that the predictors and characteristics of BPD overlap with those of hormone sensitivity. First, stress exposure and vulnerability are relevant to the etiology of BPD (Lynam & Widiger 2001; Ball & Links 2009) and associated with hormone sensitivity (Gollenberg *et al*. 2010; Gordon *et al*. 2015; Nillni *et al*. 2015; Eisenlohr-Moul *et al*. 2016). Second, impulsivity, a core feature of BPD (Peters *et al*. 2013), predicts sensitivity to hormone withdrawal in rodents (Löfgren *et al*. 2009) and humans (Martel *et al*. 2017). Finally, *behavioral* manifestations of emotion-related impulsivity show PME, including binge eating in women with bulimia nervosa and non-clinical samples of women from the community (Edler *et al*. 2007; Klump *et al*. 2013), substance use and abuse (Terner & de Wit 2006), and suicide attempts (Saunders & Hawton 2006).

### Clinical Implications

Despite development of useful psychotherapies for BPD, such as dialectical behavior therapy (DBT; Linehan 2014) and mentalization-based therapy (Bateman & Fonagy 2004), quality of life often remains low following treatment (Linehan *et al*. 2006). Experimental work is needed to determine whether stabilization of ovarian steroid flux may relieve PME of BPD. Greater cycle awareness in the context of psychosocial treatment may support targeted skills use to improve symptom stability (e.g., integration of menses into a DBT diary card). Finally, the dramatic PME of hopelessness observed here combines with previous work (Saunders & Hawton 2006) to highlight cyclical changes in suicide risk.

### Limitations and Strengths

This work has important limitations. The clinical sample was small; although repeated measures mitigated power concerns somewhat, results need replication. There was no clinical or healthy control group; although use of the C-PASS (Eisenlohr-Moul *et al*. 2017b) ensures that the observed PMEs were clinically significant, the cyclical changes observed here may be due to transdiagnostic factors (e.g., trauma exposure; Eisenlohr-Moul *et al*. 2016) rather than being specific to BPD. Women were excluded for OCs; since some people discontinue OCs following adverse reactions, those not taking OCs could be more likely to exhibit hormone sensitivity. Unfortunately, salivary E2 measurements did not meet quality control standards; future work may benefit from morning sampling and expediting transfer of samples to lower-temperature storage. Further, salivary steroid sampling was too infrequent for adequately-powered hormone analyses; future work should utilize more frequent sampling and eventually use experimental designs.

This work also has notable strengths relative to past work. Study purpose was blinded during recruitment to reduce self-selection issues. Participants were unmedicated, community-recruited, and demographically diverse. Use of online (rather than paper) surveys ensured proximal, time-verified symptom ratings. Both LH surge tests and salivary P4 were used to confirm ovulation and cycle phase. Finally, we examined all pairwise phase comparisons to precisely describe cyclical symptom change in BPD.

### Conclusions

People with BPD suffer intense fluctuations in painful emotional and behavioral symptoms. The present work supports the hypothesis that females with BPD may be at high risk of PME, with secondary worsening in the midluteal (high arousal symptoms) and follicular (low arousal symptoms) phases. Patterns of symptom change suggest shared pathophysiology with PMDD with regard to midluteal and perimenstrual high-arousal symptoms, but also suggest potentially divergent pathophysiology with regard to depression. Experimental studies are needed for the dual purposes of investigating the therapeutic potential of hormone stabilization while probing the pathophysiologic relevance of steroid changes to cyclical worsening of BPD.

## References

Aviram RB, Brodsky BS, Stanley B (2006). Borderline personality disorder, stigma, and treatment implications. Harvard review of psychiatry 14, 249–256.

Baer RA, Sauer SE (2011). Relationships between depressive rumination, anger rumination, and borderline personality features. Personality disorders 2, 142–150.

Ball JS, Links PS (2009). Borderline personality disorder and childhood trauma: evidence for a causal relationship. Springer Current psychiatry reports 11, 63–68.

Bateman AW, Fonagy P (2004). Mentalization-based treatment of BPD. Journal of personality disorders 18, 36–51.

Bender DS, Dolan RT, Skodol AE, Sanislow CA, Dyck IR, McGlashan TH, Shea MT, Zanarini MC, Oldham JM, Gunderson JG (2001). Treatment utilization by patients with personality disorders. The American journal of psychiatry 158, 295–302.

Ben Dor R, Harsh VL, Fortinsky P, Koziol DE, Rubinow DR, Schmidt PJ (2013). Effects of pharmacologically induced hypogonadism on mood and behavior in healthy young women. The American journal of psychiatry 170, 426–433.

Bernstein DP, Fink L (1998). Childhood trauma questionnaire: A retrospective self-report: Manual. Psychological Corporation.

Bloch M, Schmidt PJ, Danaceau M, Murphy J, Nieman L, Rubinow DR (2000). Effects of gonadal steroids in women with a history of postpartum depression. The American journal of psychiatry 157, 924–930.

Dawson DN, Eisenlohr-Moul TA, Paulson JL, Peters JR, Rubinow DR, Girdler SS (n.d.). Emotion-related impulsivity and rumination predict the perimenstrual severity and trajectory of symptoms in women with a menstrually related mood disorder. Journal of clinical psychology

Edler C, Lipson SF, Keel PK (2007). Ovarian hormones and binge eating in bulimia nervosa. Psychological medicine 37, 131–141.

Eisenlohr-Moul TA, DeWall CN, Girdler SS, Segerstrom SC (2015). Ovarian hormones and borderline personality disorder features: Preliminary evidence for interactive effects of estradiol and progesterone. Biological psychology 109, 37–52.

Eisenlohr-Moul TA, Girdler SS, Johnson JL, Schmidt PJ, Rubinow DR (2017a). Treatment of premenstrual dysphoria with continuous versus intermittent dosing of oral contraceptives: Results of a three-arm randomized controlled trial. Depression and anxiety. In Press.

Eisenlohr-Moul TA, Girdler SS, Schmalenberger KM, Dawson DN, Surana P, Johnson JL, Rubinow DR (2017b). Toward the Reliable Diagnosis of DSM-5 Premenstrual Dysphoric Disorder: The Carolina Premenstrual Assessment Scoring System (C-PASS). The American journal of psychiatry 174, 51–59.

Eisenlohr-Moul TA, Rubinow DR, Schiller CE, Johnson JL, Leserman J, Girdler SS (2016). Histories of abuse predict stronger within-person covariation of ovarian steroids and mood symptoms in women with menstrually related mood disorder. Psychoneuroendocrinology 67, 142–152.

Endicott J, Nee J, Harrison W (2006). Daily Record of Severity of Problems (DRSP): reliability and validity. Archives of women’s mental health 9, 41–49.

Epperson CN, Steiner M, Hartlage SA, Eriksson E, Schmidt PJ, Jones I, Yonkers KA (2012). Premenstrual dysphoric disorder: evidence for a new category for DSM-5. The American journal of psychiatry 169, 465–475.

First MB, Gibbon M, Spitzer RL, Benjamin LS, Williams JBW (1997). Structured Clinical Interview for DSM-IV Axis II Personality Disorders: SCID-II. American Psychiatric Press.

Freeman EW, Halberstadt SM, Rickels K, Legler JM, Lin H, Sammel MD (2011). Core symptoms that discriminate premenstrual syndrome. Journal of women’s health 20, 29–35.

Glenn CR, Cha CB, Kleiman EM, Nock MK (2017). Understanding Suicide Risk within the Research Domain Criteria (RDoC) Framework: Insights, Challenges, and Future Research Considerations. Clinical psychological science 5, 568–592.

Gollenberg AL, Hediger ML, Mumford SL, Whitcomb BW, Hovey KM, Wactawski-Wende J, Schisterman EF (2010). Perceived stress and severity of perimenstrual symptoms: the BioCycle Study. Journal of women’s health 19, 959–967.

Gordon JL, Girdler SS, Meltzer-Brody SE, Stika CS, Thurston RC, Clark CT, Prairie BA, Moses-Kolko E, Joffe H, Wisner KL (2015). Ovarian hormone fluctuation, neurosteroids, and HPA axis dysregulation in perimenopausal depression: a novel heuristic model. The American journal of psychiatry 172, 227–236.

Gordon JL, Rubinow DR, Eisenlohr-Moul TA, Leserman J, Girdler SS (2016). Estradiol variability, stressful life events, and the emergence of depressive symptomatology during the menopausal transition. Menopause 23, 257–266.

Halbreich U, Endicott J, Schacht S, Nee J (1982). The diversity of premenstrual changes as reflected in the Premenstrual Assessment Form. Acta psychiatrica Scandinavica 65, 46–65.

Hammarbäck S, Ekholm UB, Bäckström T (1991). Spontaneous anovulation causing disappearance of cyclical symptoms in women with the premenstrual syndrome. Acta endocrinologica 125, 132–137.

Hartlage SA, Brandenburg DL, Kravitz HM (2004). Premenstrual exacerbation of depressive disorders in a community-based sample in the United States. Psychosomatic medicine 66, 698–706.

Hengartner MP, Kruger THC, Geraedts K, Tronci E, Mancini T, Ille F, Egli M, Röblitz S, Ehrig R, Saleh L, Spanaus K, Schippert C, Zhang Y, Leeners B (2017). Negative affect is unrelated to fluctuations in hormone levels across the menstrual cycle: Evidence from a multisite observational study across two successive cycles. Journal of psychosomatic research 99, 21–27.

Hyler SE, Skodol AE, Oldham JM, Kellman HD, Doidge N (1992). Validity of the Personality Diagnostic Questionnaire-Revised: a replication in an outpatient sample. Comprehensive psychiatry 33, 73–77.

Klump KL, Keel PK, Racine SE, Burt SA, Burt AS, Neale M, Sisk CL, Boker S, Hu JY (2013). The interactive effects of estrogen and progesterone on changes in emotional eating across the menstrual cycle. Journal of abnormal psychology 122, 131–137.

Leeners B, Kruger THC, Geraedts K, Tronci E, Mancini T, Ille F, Egli M, Röblitz S, Saleh L, Spanaus K, Schippert C, Zhang Y, Hengartner MP (2017). Lack of Associations between Female Hormone Levels and Visuospatial Working Memory, Divided Attention and Cognitive Bias across Two Consecutive Menstrual Cycles. Frontiers in behavioral neuroscience 11, 120.

Linehan M (2014). DBT Skills Training Manual. Guilford Publications.

Linehan MM, Comtois KA, Murray AM (2006). Two-year randomized controlled trial and follow-up of dialectical behavior therapy vs therapy by experts for suicidal behaviors and borderline personality …. jamanetwork.com Archives of general psychiatry

Löfgren M, Johansson I-M, Meyerson B, Turkmen S, Bäckström T (2009). Withdrawal effects from progesterone and estradiol relate to individual risk-taking and explorative behavior in female rats. Physiology & behavior 96, 91–97.

Lynam DR, Widiger TA (2001). Using the five-factor model to represent the DSM-IV personality disorders: an expert consensus approach. Journal of abnormal psychology 110, 401–412.

Marschall D, Sanftner J, Tangney JP (1994). The state shame and guilt scale. Fairfax, VA: George Mason University

Martinez PE, Rubinow DR, Nieman LK, Koziol DE, Leslie Morrow A, Schiller CE, Cintron D, Thompson KD, Khine KK, Schmidt PJ (2013). 5α-Reductase Inhibition Prevents the Luteal Phase Increase in Plasma Allopregnanolone Levels and Mitigates Symptoms in Women with Premenstrual Dysphoric Disorder. Neuropsychopharmacology: official publication of the American College of Neuropsychopharmacology 41, 1093–1102.

Morey LC (2014). Personality Assessment Inventory (PAI). In The Encyclopedia of Clinical Psychology. John Wiley & Sons, Inc.

Nillni YI, Pineles SL, Patton SC, Rouse MH, Sawyer AT, Rasmusson AM (2015). Menstrual cycle effects on psychological symptoms in women with PTSD. Journal of traumatic stress 28, 1–7.

Paris J, Zweig-Frank H (2001). A 27-year follow-up of patients with borderline personality disorder. Comprehensive psychiatry 42, 482–487.

Peters JR, Eisenlohr-Moul TA, Upton BT, Talavera NA, Folsom JJ, Baer RA (2017). Characteristics of Repetitive Thought Associated with Borderline Personality Features: A Multimodal Investigation of Ruminative Content and Style. Journal of psychopathology and behavioral assessment, 1–11.

Peters JR, Upton BT, Baer RA (2013). Brief report: relationships between facets of impulsivity and borderline personality features. Journal of personality disorders 27, 547–552.

Putnam KT, Wilcox M, Robertson-Blackmore E, Sharkey K, Bergink V, Munk-Olsen T, Deligiannidis KM, Payne J, Altemus M, Newport J, Apter G, Devouche E, Viktorin A, Magnusson P, Penninx B, Buist A, Bilszta J, O’Hara M, Stuart S, Brock R, Roza S, Tiemeier H, Guille C, Epperson CN, Kim D, Schmidt P, Martinez P, Di Florio A, Wisner KL, Stowe Z, Jones I, Sullivan PF, Rubinow D, Wildenhaus K, Meltzer-Brody S, Postpartum Depression: Action Towards Causes and Treatment (PACT) Consortium (2017). Clinical phenotypes of perinatal depression and time of symptom onset: analysis of data from an international consortium. The Lacet Psychiatry 4, 477–485.

Saunders KEA, Hawton K (2006). Suicidal behaviour and the menstrual cycle. Psychological medicine 36, 901–912.

Schiller CE, Johnson SL, Abate AC, Schmidt PJ, Rubinow DR (2016). Reproductive Steroid Regulation of Mood and Behavior. Comprehensive Physiology 6, 1135–1160.

Schiller CE, Schmidt PJ, Rubinow DR (2014). Allopregnanolone as a mediator of affective switching in reproductive mood disorders. Psychopharmacology 231, 3557–3567.

Schmidt PJ, Ben Dor R, Martinez PE, Guerrieri GM, Harsh VL, Thompson K, Koziol DE, Nieman LK, Rubinow DR (2015). Effects of Estradiol Withdrawal on Mood in Women With Past Perimenopausal Depression: A Randomized Clinical Trial. JAMA psychiatry 72, 714–726.

Schmidt PJ, Keenan PA, Schenkel LA, Berlin K, Gibson C, Rubinow DR (2013). Cognitive performance in healthy women during induced hypogonadism and ovarian steroid addback. Archives of women’s mental health 16, 47–58.

Schmidt PJ, Nieman LK, Danaceau MA, Adams LF, Rubinow DR (1998). Differential behavioral effects of gonadal steroids in women with and in those without premenstrual syndrome. The New England journal of medicine 338, 209–216.

Schwartz DH, Romans SE, Meiyappan S, De Souza MJ, Einstein G (2012). The role of ovarian steroid hormones in mood. Hormones and behavior 62, 448–454.

Shirtcliff EA, Granger DA, Schwartz EB, Curran MJ, Booth A, Overman WH (2000). Assessing estradiol in biobehavioral studies using saliva and blood spots: simple radioimmunoassay protocols, reliability, and comparative validity. Hormones and behavior 38, 137–147.

Smith SS, Ruderman Y, Frye C, Homanics G, Yuan M (2006). Steroid withdrawal in the mouse results in anxiogenic effects of 3α,5β-THP: a possible model of premenstrual dysphoric disorder. Psychopharmacology 186, 323–333.

Snijders TAB (2005). Power and Sample Size in Multilevel Linear Models. In Encyclopedia of Statistics in Behavioral Science. John Wiley & Sons, Ltd.

Spielberger CD (1983). Manual for the State-Trait Anxiety Inventory STAI.

Stein MB, Schmidt PJ, Rubinow DR, Uhde TW (1989). Panic disorder and the menstrual cycle: panic disorder patients, healthy control subjects, and patients with premenstrual syndrome. The American journal of psychiatry 146, 1299–1303.

Stoffel EC, Craft RM (2004). Ovarian hormone withdrawal-induced “depression” in female rats. Physiology & behavior 83, 505–513.

Sukhodolsky DG, Golub A, Cromwell EN (2001). Development and validation of the anger rumination scale. Personality and individual differences 31, 689–700.

Terner JM, de Wit H (2006). Menstrual cycle phase and responses to drugs of abuse in humans. Drug and alcohol dependence 84, 1–13.

Ziv B, Russ MJ, Moline M, Hurt S, Zendell S (1995). Menstrual Cycle Influences On Mood and Behavior in Women With Borderline Personality Disorder. Journal of personality disorders 9, 68–75.

